# Chromosome Integrity is Required for the Initiation of Meiotic Sex Chromosome Inactivation in *Caenorhabditis elegans*

**DOI:** 10.1101/2020.11.05.369132

**Authors:** Yisrael Rappaport, Hanna Achache, Roni Falk, Omer Murik, Oren Ram, Yonatan B. Tzur

**Affiliations:** Department of Genetics, The Institute of Life Sciences, The Hebrew University of Jerusalem, Jerusalem 91904, Israel; Medical Genetics Institute, Shaare Zedek Medical Center, Jerusalem, Israel; Department of Biological Chemistry, The Institute of Life Sciences, The Hebrew University of Jerusalem, Jerusalem, Israel

**Keywords:** Meiosis, MSCI, meiotic sex chromosome inactivation, germline

## Abstract

During meiosis of heterogametic cells, such as XY meiocytes, sex chromosomes of many species undergo transcriptional silencing known as meiotic sex chromosome inactivation (MSCI). Silencing also occurs in aberrantly unsynapsed autosomal chromatin. The silencing of unsynapsed chromatin, is assumed to be the underline mechanism for MSCI. Initiation of MSCI is disrupted in meiocytes with sex chromosome-autosome translocations. Whether this is due to aberrant synapsis or the lack of sex chromosome integrity has never been determined. To address this, we used CRISPR to engineer *Caenorhabditis elegans* stable strains with broken X chromosomes that didn’t undergo translocations with autosomes. In early meiotic nuclei of these mutants, the X fragments lack silent chromatin modifications and instead the fragments are enriched with transcribing chromatin modifications. Moreover, the level of active RNA polymerase II staining on the X fragments in mutant nuclei is similar to that on autosomes, indicating active transcription on the X. Contrary to previous models, which predicted that any unsynapsed chromatin is silenced during meiosis, X fragments that did not synapse were robustly stained with RNA polymerase II and gene expression levels were high throughout the broken X. Therefore, lack of synapsis does not trigger MSCI if sex chromosome integrity is lost. Moreover, our results suggest that a unique character of the chromatin of sex chromosomes underlies their lack of meiotic silencing due to both unsynapsed chromatin and sex chromosome mechanisms when their integrity is lost.

---

During prophase I of meiosis in most sexually reproducing organisms, homologous chromosomes pair and then undergo a closer engagement known as synapsis to complete interhomolog crossover recombination ^1–12^. In the heterogametic cells of many species (e.g., meiocytes with X and Y chromosomes), the sex chromosomes pair but undergo synapsis and crossovers only in the pseudo homology regions. In mouse testes, these paired chromosomes form a compartment of heterochromatic chromatin referred to as the XY body, which undergoes transcriptional silencing through many stages of meiosis and, in some cases, into gametogenesis. Although sex chromosomes have appeared and disappeared several times during metazoan evolution, meiotic sex chromosomes inactivation (MSCI) occurs in many species from worms to humans ^13–17^.

MSCI in mammals is perturbed by mutations in genes involved in meiotic double-strand break formation (e.g., *Spo11*), DNA damage response (e.g., *Brca1, Mdc1, Topbp1*, *and Setx* ^18–22^), and chromatin modifiers (e.g., S*etdb1* ^23^). In mouse testes, the lack of MSCI usually leads to pachytene arrest, apoptosis, and persistence of homologous recombination intermediates ^24, 25^. Although the molecular mechanism of MSCI emplacement is well characterized, our knowledge of how MSCI is triggered is lacking.

In the nematode *C. elegans,* MSCI is present in both XO male and XX hermaphrodite worms. In gonads of adult worms, nuclei are arranged according to developmental progression. At the distal end, proliferative cells undergo mitotic divisions, and they enter meiosis at the leptotene/zygotene stage, where homologous chromosomes pair. Pairing is closely followed by synapsis within an evolutionary conserved structure involving lateral and central proteinaceous elements that keep the homologs aligned. The chromosomes are fully synapsed during pachytene, which allows crossovers to mature. In hermaphrodite worms, the nuclei proceed through diplotene and reach maturity at the diakinesis stage ^7, 8, 12^. In male worms, the single X chromosome does not undergo synapsis and is transcriptionally silenced throughout meiosis ^26–29^. In hermaphrodites, the two X chromosomes pair and synapse, yet these chromosomes are silenced in early meiotic stages; toward the end of pachytene the silencing is relieved, however, and transcription from these chromosomes increases.

The current model views MSCI as a special case of meiotic silencing of unsynapsed chromatin (MSUC) ^30^, a processes characterized in mammals, *Neurospora crassa*, and *C. elegans* ^30–32^. Several lines of evidence support this model, including the silencing of the unsynapsed X chromosome in XO female mouse meiocytes ^30^ and the lack of silencing in synapsed Y chromosomes in mouse XYY testes ^33^. Furthermore, when translocations between autosomes and sex chromosomes occur, the localization of MSCI effectors to the sex chromosomes fragments is perturbed ^34–37^. This lack of MSCI was explained by the aberrant synapsis of the sex chromosome fragments observed in these nuclei. Nevertheless, several reports suggest that, in some cases, synapsed translocated sex chromosomes show MSCI markers ^34–37^ raising the possibility that changes in sex chromosome integrity can perturb MSCI.

In this study we tested the hypothesis that sex chromosomes must be unbroken (hence chromosome integrity) for efficient MSCI. We created stable worm strains with broken X chromosomes that did not translocate to autosomes. We found that in meiocytes of these strains, and in a strain with a reciprocal translocation of chromosomes V and X, MSCI failed to initiate. The X chromosome segments showed active transcription markers, and the expression of X-linked genes in the gonads was increased in the strains with broken X chromosomes. In contrast to the prediction that MSCI is a special case of MSUC, we showed that segments of the X that are unsynapsed are not silenced. Loss of MSCI was accompanied by meiotic defects, perturbations in DNA repair, and reduced fertility. Based on these data, we suggest that chromosome integrity is required in *C. elegans* hermaphrodites for MSCI and proper meiotic progression.

## Results

### Creation of *C. elegans* stable homozygous strains with broken X chromosomes

Previous reports indicated that MSCI is disrupted in heterogametic cells with sex-chromosome to autosome translocations and that, in some cases, gene expression was uncoupled from the synapsis state of the translocated chromosomes ^30, 34, 35, 37, 38^. This suggested that disruption of chromosome integrity prevents initiation of MSCI. To test this hypothesis, we aimed to create worm strains with an X chromosome with disrupted integrity but without a translocation with an autosome, reducing the possibility of aberrant synapsis. Ideally, we wanted a system that 1) is homozygous stable, 2) has fragments considerably smaller than the full-size chromosome but larger than extra-chromosomal arrays and free duplications, and 3) has fragments with telomeres on both sides.

Previous reports indicate that multiple CRISPR-mediated DNA double-strand breaks at homologous chromosomal loci can lead to chromosomal aberrations such as inversions, large deletions, circularizations, and chromosomal cleavages ^39–45^. To create strains with fragmented X chromosomes, we searched for genomic regions near the ends of chromosome X with homology to regions at the center. If breaks at both at both center and one of the ends loci are formed, and not repaired, three fragments are created. If two non-adjacent breaks are ligated, two fragments result. If all three fragments are ligated, then chromosome rearrangements may occur. A fragment without telomers could also undergo circularization as was detected before ^39, 46^. We identified a 2.2-kb region (X:16508962-16511217) on the right side of the X chromosome encompassing the non-coding gene *linc-20*, which is homologous (>92% identity) to a region near the center of the X chromosome (X:7769295-7771552) within the fourteenth intron (i14) of *deg-1*. Neither of these genes have previously been associated with germline roles ^47–49^. As previously described ^50, 51^, we directed Cas9 to these loci with four guide RNAs (gRNAs) to create multiple breaks. We assayed the progeny of injected worms for deletions in targeted loci using PCR and isolated a strain with small deletions in both: The deletion in i14 was 2597 bases, and two deletions were observed in *linc-20* of 1417 and 2721 bases (data not shown). After five outcrosses with the wild-type strain, the YBT7 strain was established. All further experiments were conducted using this outcrossed strain. This strain was maintained through multiple generations without any change in genotyping markers of these loci.

We next evaluated whether there are structural alterations in the X chromosomes of YBT7 worms using Nanopore long-read DNA sequencing. This analysis indicated that Cas9-mediated cleavages in i14 of *deg-1* and in *linc-20* loci resulted in fusion of the internal fragment from X:772344 to X:16511091 into a circular chromosome of approximately 8.7 Mbp. Additionally, the left fragment was ligated to the right fragment (linking X:7769697 to X:16513803), creating an approximately 9-Mbp linear chromosome (Fig. 1a and supplemental data). We also detected a small inversion within the fusion point of the linear chromosome (X:7762996 to X:16513802). Sanger sequencing confirmed the fusion points of these fragments. No other major chromosomal alterations were detected by the Nanopore sequencing.

**Figure 1.**
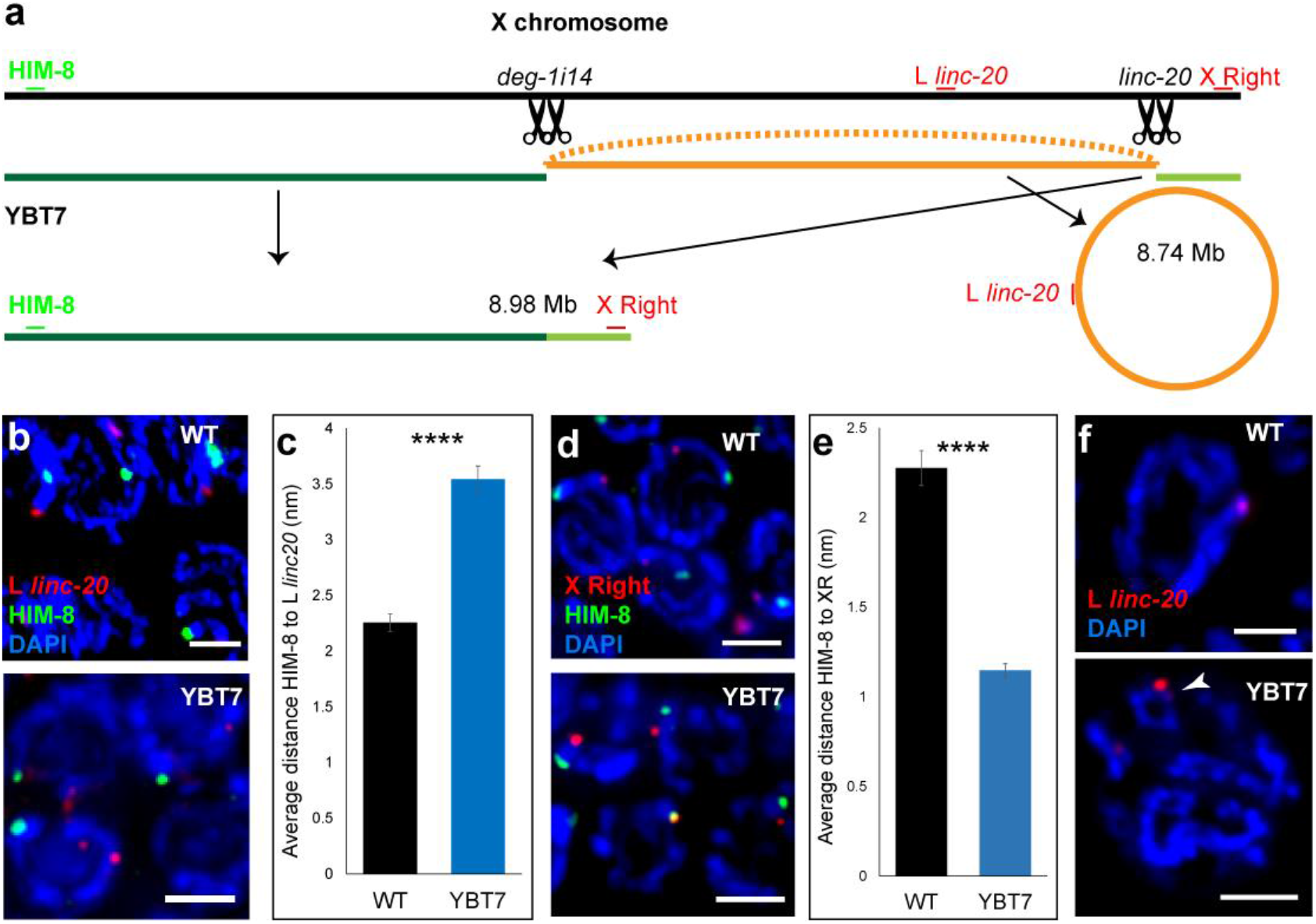
Engineering of worm strains with broken X chromosomes. **a,** Illustration of the X chromosome in a wild-type worm and the fragments resulting from Cas9-mediated cleavage in YBT7 worms. The gRNA binding sites (black scissors) and cytological markers (green and red) used in analyses of X chromosome fragmentation are indicated. **b,** Pachytene nuclei stained with DAPI (blue), HIM-8 (green), and left *linc-20* FISH probe (red). **c,** Quantification of the distance between HIM-8 and the site left of *linc-20*. **d,** Pachytene nuclei stained with DAPI (blue), HIM-8 (green), and FISH probe complementary to the right end of the X chromosome (red). **e,** Quantification of the distance between HIM-8 and the site on the right end of chromosome X. **f,** Late pachytene nuclei stained with DAPI (blue) and the FISH probe marking a site left of *linc-20* (red). FISH signal associated with a circular chromosome is marked with an arrowhead. n≥80. **** *p*<0.0001, Mann-Whitney test. Scale bars = 3 μM.

We verified that both the X chromosomes are fragmented in YBT7 by co-staining YBT7 gonads with antibodies against HIM-8, a protein that binds the left end of the X chromosome ^52^, and with fluorescent in situ hybridization (FISH) probes directed to a site left of *linc-20* locus (L *linc-20*). In YBT7, HIM-8 is predicted to bind the linear fragment, whereas the FISH probes bind to the circular fragment (Fig. 1a). In wild-type pachytene nuclei these markers appeared on the same DAPI-stained track, but in YBT7 the HIM-8 and FISH staining mostly marked different DAPI tracks (Fig. 1b; 80/80 vs. 10/80 on the same track, respectively). Due to the spatial resolution of our fluorescent microscopy two very close tracks are not always differentiated, which is likely why the two markers scored on the same track in a fraction of YBT7 nuclei examined. The distance between the markers was also shorter in wild-type worms than in YBT7 worms (Fig. 1c; 2.25±0.08 μM vs. 3.5±0.1 μM, respectively, n=80). We next co-stained YBT7 gonads with HIM-8 antibodies and FISH probes directed to the right side of the chromosome (Fig. 1a). The two markers were on the same DAPI-stained track during pachytene in both strains, but in YBT7 they were closer than in the wild-type strain (Fig. 1d-e; 1.1±0.04 μM vs. 2.3±0.1 μM, respectively, n=80), suggesting that the left end of the X is closer to the right end in YBT7 than in the wild-type strain.

Circular chromosomes and large extrachromosomal circular DNA are observed in many organisms in normal and tumor cells, and circular chromosomes can be maintained through multiple mitotic divisions ^53^. In humans, these chromosomal aberrations are thought to result from two double-stranded breaks ^46^. To verify that the middle segment of the X chromosome exists as a circle in the YBT7 germline cells, we imaged late pachytene nuclei marked with the FISH expected to be within the circular fragment (Fig. 1a, the probe to the left of the *linc-20* locus). In YBT7 but not in wild-type gonads, we detected nuclei in which this probe was localized to a circular DAPI stained track (Fig. 1f). Taken together, these analyses indicate that YBT7 worm cells have a stably fragmented X chromosome. These worms are homozygous for two dissociated parts of the X chromosome that are not translocated to autosomes.

### Loss of MSCI markers in early meiotic nuclei with broken X chromosome

In the gonads of hermaphroditic *C. elegans*, the two X chromosomes are silenced from the proliferative nuclei until late pachytene, and then transcription resumes ^26–29, 54^. During early meiotic stages, the chromatin of X chromosomes is enriched with modifications correlated with low transcriptional activity such as histone H3 trimethylated at lysine 27 (H3K27me3) ^55^. We tested whether the disruption of X chromosome integrity changed the chromatin state by staining the gonads with H3K27me3 antibodies and with HIM-8 to mark the X chromosome. As shown previously ^55^, we found that in early wild-type pachytene nuclei the X chromosomes were strongly stained with H3K27me3 antibodies (Fig. 2a). In YBT7 early pachytene nuclei, the HIM-8 marked DAPI track was stained with the H3K27me3 antibody at levels similar to all other DAPI tracks, and no chromosome was strongly stained (Fig. 2a). We next measured the level of H3K27me3 signal associated with the HIM-8-marked chromosome relative to the level associated with the autosomes in the same nucleus. We found that in wild-type strain the ratio was 1.5±0.07, whereas that in the YBT7 strain was 1.06±0.03 (Fig. 2b; n≥10, *p*<0.01 by the Mann-Whitney test). Thus, the linear fragment of the X chromosome in early meiotic YBT7 nuclei was marked by H3K27me3 at levels very similar to autosomes.

**Figure 2:**
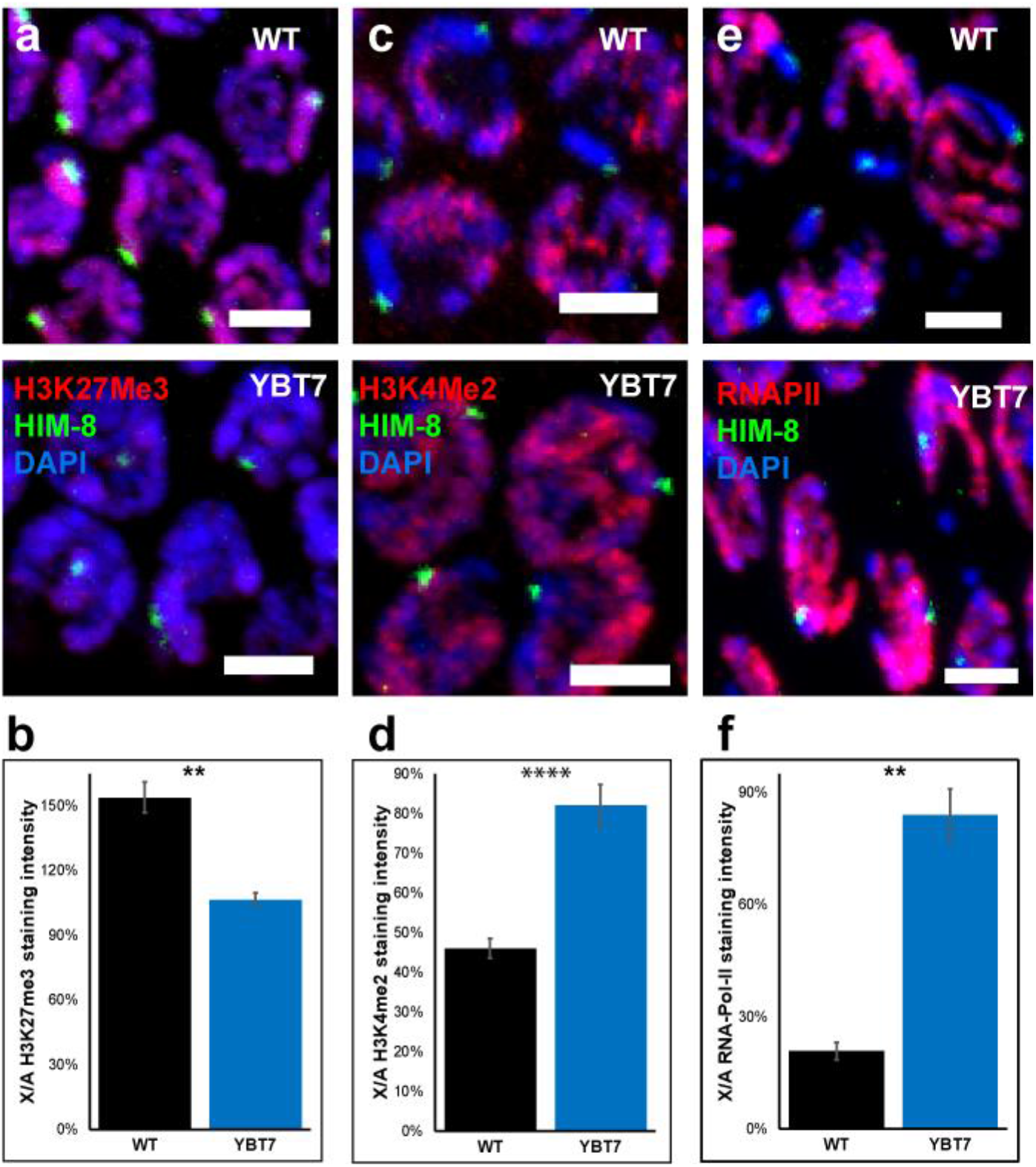
Active transcription marks are associated with the left fragment of the X chromosome in YBT7. **a,** Early pachytene nuclei stained with DAPI (blue), HIM-8 (green), and antibody against H3K27me3 (red). **b,** Average of the relative H3K27me3 signal per wild-type and YBT7 nucleus on a HIM-8-marked body vs. an autosome. **c,** Early pachytene nuclei stained with DAPI (blue), HIM-8 (green), and antibody against H3K4me2 (red). **d,** Average of the relative H3K4me2 signal per wild-type and YBT7 nucleus on a HIM-8-marked body vs. an autosome. **e,** Early pachytene nuclei stained with DAPI (blue), HIM-8 (green), and antibody against RNAPII. **f,** Average of the relative RNAPII signal per wild-type and YBT7 nucleus on a HIM-8-marked body vs. an autosome. n≥10. ** *p*<0.01, **** *p*<0.0001, Mann-Whitney test. Scale bar = 3 μM.

We next tested whether transcription from the X chromosome changes when its integrality is disrupted. For this analysis, we used antibodies to histone H3 dimethylated at lysine 4 (H3K4me2), a modification correlated with active transcription in *C. elegans*. In nuclei in early meiotic stages in wild-type gonads, there are very low levels of H3K4me2 on the X chromosomes ^26, 55, 56^. In YBT7 nuclei, however, the staining of the X chromosome fragment was more strongly stained than in wild-type gonads; the level was similar to that of autosomes (Fig. 2c). Quantification of the staining levels on the HIM-8-marked chromosome vs. the autosomes within the same nucleus indicated that the ratio was significantly higher in YBT7 nuclei than in wild-type nuclei (Fig. 2d; 0.82±0.05 vs. 0.46±0.02, respectively, n≥15,).

One of the most direct cytology markers of active transcription is the antibody that recognizes the B1 subunit of RNA polymerase II (RNAPII) when phosphorylated at Ser2 ^26, 57, 58^. As was observed previously ^26^, we found that in early pachytene nuclei of wild-type gonads the X chromosome as not strongly associated with this antibody (Fig. 2e). In contrast, in YBT7 early pachytene nuclei the chromatin tracks with the HIM-8 mark indicative of the X chromosome fragment were strongly stained for active RNAPII (Fig. 2e). Quantification of the ratio of RNAPII signal on the HIM-8 associated chromosome vs. an autosome within the same nucleus showed a dramatic difference between wild-type and YBT7 gonads (Fig. 2f; 0.2±0.02 vs. 0.8±0.07, respectively, n≥11). Taken together these results indicate that in YBT7 nuclei, the linear fragment of the broken X chromosome is not silenced during early meiotic steps.

### Many X linked genes are upregulated in YBT7 gonads

Our cytological data suggest that the X chromosome linear fragment in YBT7 gonads does not undergo meiotic silencing. To determine whether the circular fragment or specific regions of the linear fragment are transcriptionally silent, we dissected gonads from wild-type and YBT7 worms and compared their transcriptomes. We found that 197 genes out of 2867 from the X chromosome were highly upregulated and 24 were highly downregulated in YBT7 compared to wild-type gonads (Table S1). Of the highly upregulated genes, 91 are encoded on the circular fragment and 106 on the linear fragment, and no specific regions were over- or under-represented (Fig. 3, Table S1). This is probably an underestimation of the level of upregulated genes since loss of MSCI in hermaphrodites is expected to affect expression only at the distal part of the gonads, whereas we sequenced RNA from whole gonads. Moreover, mRNA abundance is higher at the proximal than in the distal side of the gonad^59^.

**Figure 3:**
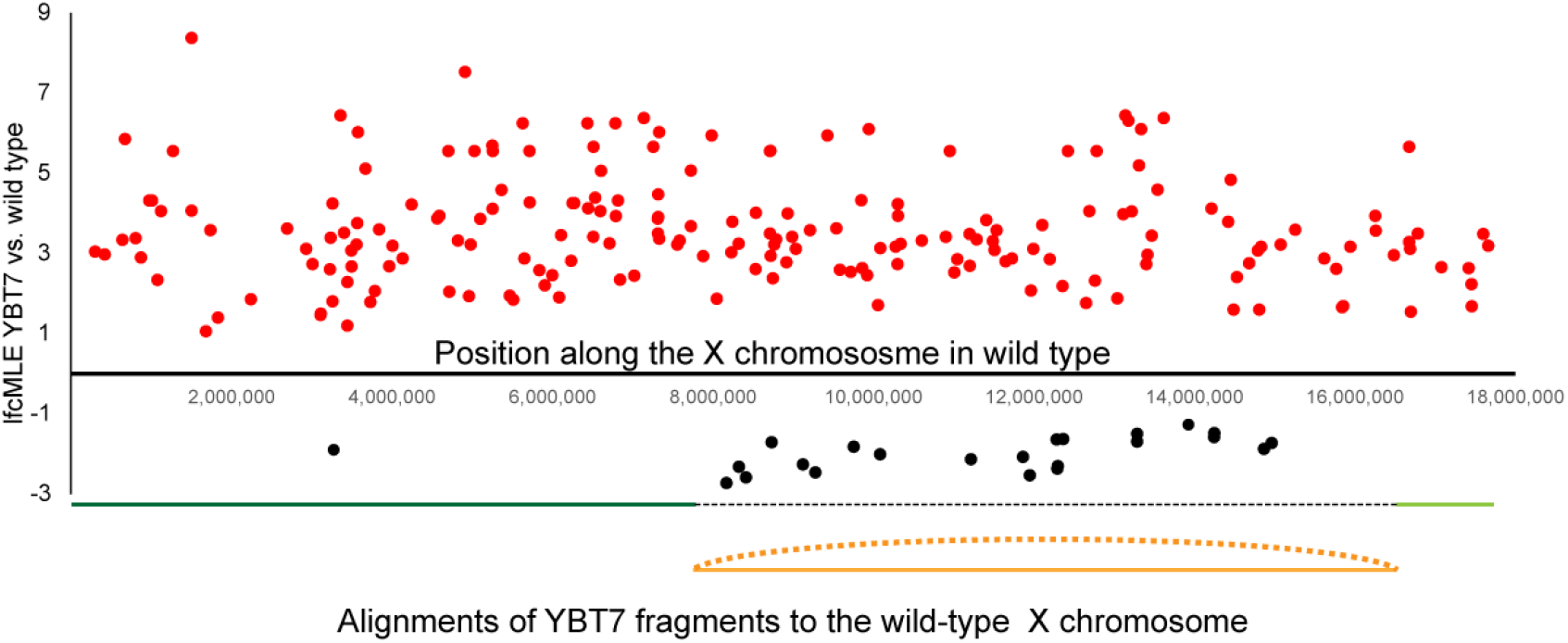
Genes along the entire X chromosome are upregulated in YBT7 gonads. Plotted is log2 of the fold-change maximum likelihood estimate (lfcMLE) of YBT7 vs. wild type for each gene along the wild-type X chromosome. Only highly differentially expressed genes are illustrated. Alignments of the linear (green) and circular (orange) fragments of YBT7 are shown schematically below the graph.

Among the 24 highly downregulated genes in YBT7 gonads, only two were on the linear fragment, and 22 on the circular fragment (Fig. 3, Table S1, *p*<0.01 by Fisher’s exact test). It is possible that one circular chromosome was lost in some meiocytes, that silencing of the circular fragment occurs only in a fraction of the meiocytes, or a complex genetic plan affecting these downregulated genes.

The dramatic difference we found in expression of genes on the X chromosome in YBT7 gonads compared to wild-type gonads could be correlated with differences in autosomal transcription. Indeed, the expression of 706 autosomal genes was also highly upregulated in YBT7 gonads (Fig. S1, Table S1). Nevertheless, A higher percentage of genes were upregulated on the fragmented X chromosome than on autosomes (*p*<7.29*10^−17^ by the hypergeometric test). The loss of silencing of X-linked genes may lead to misregulation of autosomal gene expression. However, a specific genetic plan could not be detected, suggesting a complex mechanism. Taken together these results indicate that transcription in the YBT7 germline is misregulated, and many genes from both linear and circular X chromosome fragments are more highly expressed than are the same genes in wild-type gonads.

### Worms with broken X chromosomes have severe meiotic alterations

The dramatic transcription misregulation observed in YBT7 gonads suggested that meiosis is likely disrupted in this strain. Indeed, there was a striking reduction in progeny brood size (Fig. 4a; 230±13 vs. 70±17 per worm for wild type vs. YBT7, respectively, n≥17), indicating reduced fertility. Moreover, 64±7% of the embryos laid by YBT7 worms did not hatch (Emb phenotype), whereas only 1.3±0.4 of wild-type embryos did not hatch (Fig. 4b; n≥17). This is suggestive of meiotic failure in YBT7 worms that leads to embryonic lethality. We did not detect a high incidence of males (Him phenotype), which has also been associated with failed meiotic segregations^60^.

**Figure 4:**
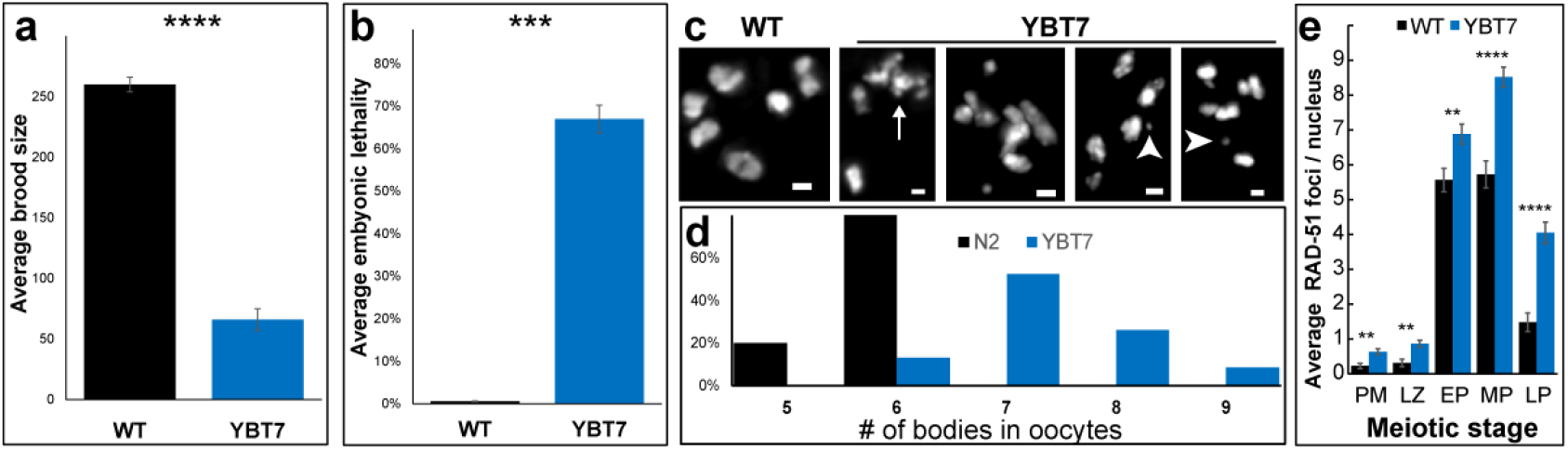
Meiotic defects are present in YBT7. **a,** Average brood and, **b,** embryonic lethality levels for wild-type and YBT7 progeny. n≥18. **c,** Mature wild-type and YBT7 oocytes stained with DAPI. Arrow indicates a chromosome aggregation. Arrowheads indicate chromosomal fragments. **d,** Percentages of DAPI stained bodies (excluding fragments) in wild-type and YBT7 oocytes. n≥23. E. Average RAD-51 foci per nucleus at different oogonial stages. n≥23. Scale bar = 1 μM. ** *p*<0.01, *** *p*<0.001, **** *p*<0.0001, Mann-Whitney test.

To obtain insight into the cellular basis of the embryonic lethality, we examined DAPI-stained wild-type and YBT7 mature oocytes. Wild-type oocytes almost always contained six DAPI-stained bodies (Fig. 4c), corresponding to the six bivalents of *C. elegans*. Many YBT7 oocytes had chromosomal aggregations, fragments, and univalent-like bodies. The number of DAPI-stained bodies varied from six to nine (excluding chromosomal fragments and aggregations, Fig. 4c-d). These chromosomal aberrations could be due to aberrant repair of DNA double-strand breaks. To test this hypothesis, we stained wild-type and YBT7 gonads with RAD-51 antibodies, which mark homologous recombination repair sites ^61–63^. In wild-type gonads, we observed previously described dynamics of RAD-51 foci ^61^: The number of foci rose during the leptotene/zygotene stage, reached a maximum during mid-pachytene, and decreased during late pachytene (Fig. 4e). In YBT7 gonads, we observed similar dynamics, but the average values in YBT7 gonads were higher in all stages (Fig. 4e). For example, in mid-pachytene we found 5.7±0.4 foci per nucleus in wild-type gonads, whereas in YBT7 gonads we found 8.5±0.3 (n≥40). Thus, double-strand break repair is perturbed in the YBT7 strain. In agreement with these results we identified that some YBT7 oocytes had very small DAPI bodies, characteristic of chromosomal fragments, as well as chromosome aggregations (Fig. 4c), which are known to be a result of aberrant DNA double strand break repair ^64, 65^. These results suggest that the X chromosome cleavage we engineered in the YBT7 strain caused perturbations in double-strand break repair and reduced fertility.

### The meiotic defects in YBT7 are not the result of the deletions in *deg-1* and *linc-20* loci

To fragment the X chromosome, we had to delete regions of the long non-coding RNA gene, *linc-20,* locus and of an intron of *deg-1*. These deletions could theoretically be the cause of the meiotic defects we observed in the YBT7 strain. To rule this out, we engineered a gene disruption in *deg-1* (*deg-1(huj28))*. This mutation did not lead to reduced brood size or embryonic lethality phenotypes (Fig. S2a-b). Similarly, a strain we engineered with a full deletion of *linc-20* had normal brood size and levels of embryonic lethality (Fig. S2c-d). These results suggest that the meiotic defects in YBT7 are not the result of the loss-of-function of either *deg-1* or *linc-20*.

Although we did not detect meiotic defects in strains with mutations in *deg-1* or *linc-20* genes, it is possible that the specific deletions we created in those sites led to the meiotic phenotypes and not the segmentation of the X chromosome. To interrogate whether the deletions are related to the phenotypes we observed, we used homology-directed repair CRISPR engineering ^66–74^ to create a strain with the three deletions within the *deg-1* and *linc-20* loci present in YBT7 but without fragmentation of the X chromosome. Brood size and embryonic lethality of this strain were equivalent to wild type (Fig. S2e-f). Taken together these results show that the phenotypes of YBT7 are not the result of deletions in the *deg-1* and *linc-20* loci.

### Aberrant MSCI occurs in another strain with a broken X chromosome

To verify that the phenotypes we observed in YBT7 stem from loss of sex chromosome integrity, we engineered another strain, YBT68, in which the X chromosome was cleaved into two fragments of roughly equal size. We isolated a strain with small deletions within i14 of *deg-1* and within the *linc-20* locus. Nanopore long read sequencing data suggested that the X chromosome is non-continuous at the *deg-1* locus, but there was no indication of fragmentation at the *linc-20* locus or of a translocation (Fig. 5a and supplementary data). The most probable interpretation of these results is that the X was broken at the *deg-1* locus with some overlap between the right and left sides of the break (Fig. 5a and supplementary data). In line with this option, staining with HIM-8 antibodies and a FISH probe directed to the right side of the X were present on the same DAPI stained track in 100% of wild type pachytene nuclei, but only in 4% of YBT68 nuclei (Fig. 5b, n≥17, *p* value<0.00001, Fisher exact test). The HIM-8 and FISH foci were spatially further from each other in YBT68 than in wild type gonads (Fig. 5b-c, 3.0±-0.1 μM vs 2.3±0.1 μM respectively, n≥17). Staining with HIM-8 and a probe directed to the left side of *linc-20* locus showed similar results (data not shown). Mature YBT68 oocytes stained with DAPI contained mostly seven bodies (Fig. 5d-e). These data indicate that in the YBT68 strain, the X chromosome is broken into two fragments.

**Figure 5:**
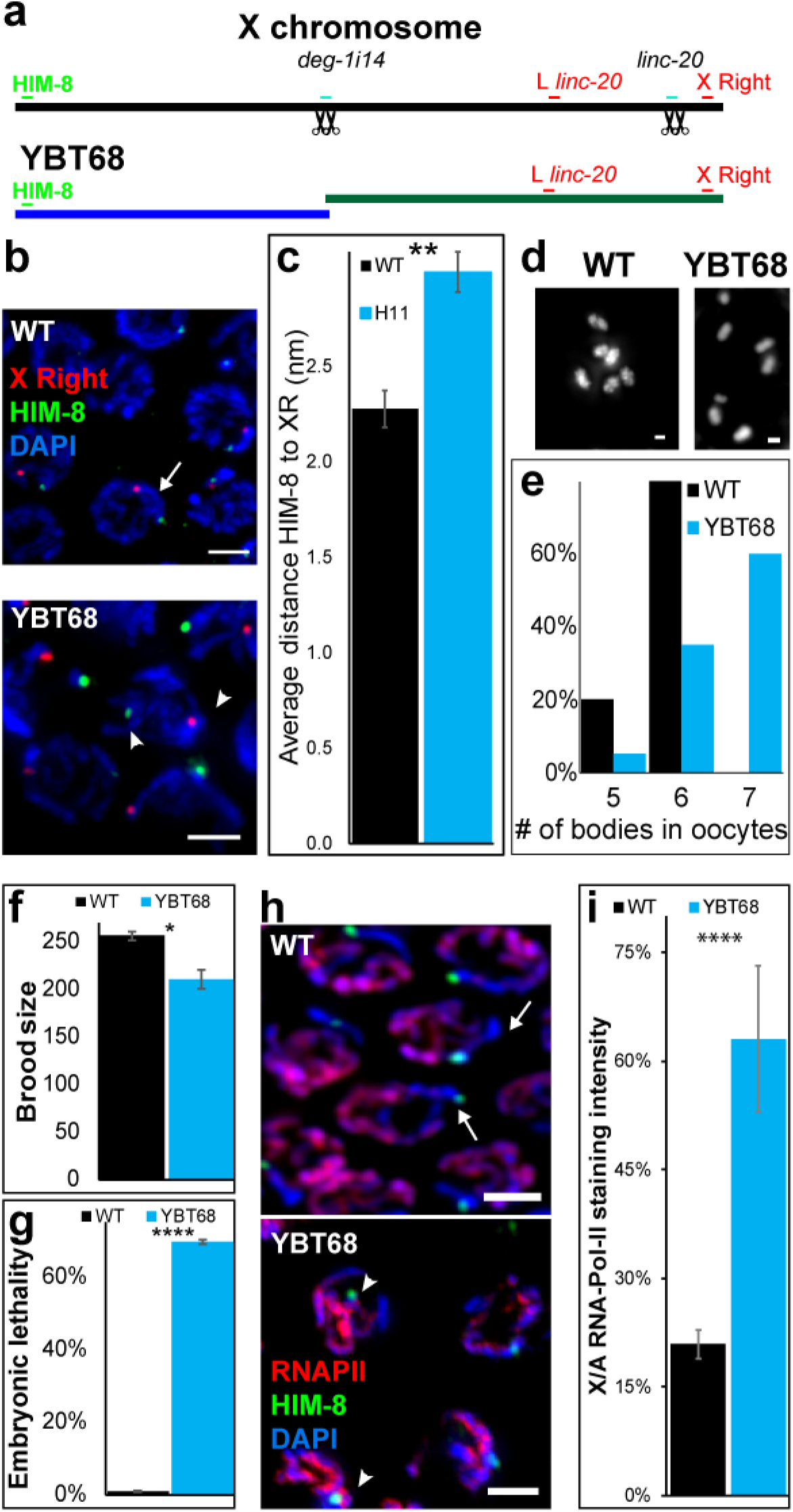
Broken X chromosomes lead to defects in MSCI and meiosis. **a,** Illustration of the X chromosomes in the wild-type and YBT68 strains. Sites of gRNAs (black scissors) and cytological markers (green and red) in the wild-type and YBT68 are marked. **b,** Pachytene nuclei stained with DAPI (blue), HIM-8 (green), and FISH probe directed to the right end of the X chromosome (red). Arrow indicates hybridization of both probes on the same chromosome. Arrowheads indicate probe hybridization on different chromosomes. Scale bar = 3 μM. **c,** Quantification of the distance between HIM-8 and the FISH probe in wild-type and YBT68 pachytene nuclei. **d,** Mature wild-type and YBT68 oocytes stained with DAPI. Scale bar = 1 μM. **e,** Distribution of the numbers of DAPI-stained bodies in wild-type and YBT68 oocytes. n≥10. **f**, Average brood size and, **g,** embryonic lethality of wild-type and YBT68 broods. **h,** Early pachytene nuclei from wild-type and YBT68 gonads stained with DAPI (blue), HIM-8 (green), and RNAPII (red). Scale bar = 3 μM. Arrows indicate HIM-8-marked chromosome with no RNAPII staining. Arrowheads indicate HIM-8-marked chromosome with significant RNAPII staining. **i,** Average of the relative RNAPII staining levels per wild-type and YBT68 nucleus on the HIM-8-marked body vs. an autosome. * *p*<0.05, ** *p*<0.01, **** *p*<0.0001, Mann-Whitney test.

We next sequenced the genomes of the parental wild-type strain, YBT7, and YBT68 at approximately 100X coverage using Illumina next-generation sequencing. The sequencing results revealed that there are no off-target structural alternations or mutations within coding genes shared between these strains (Table S2).

If MSCI initiation is dependent on X chromosome integrity, YBT68 should have similar phenotypes to YBT7. Indeed, the average brood size of YBT68 worms was significantly smaller than that of the wild-type worms (Fig. 5f), and there was over 60% embryonic lethality (Fig. 5g). We note that the progeny in YBT68 is higher than YBT7, and the nature of the chromosomal aberrations in mature oocytes is different (compare Fig. 4 to Fig. 5). This could stem from the specific chromosomal outcome of YBT7: the circular chromosome and/or no telomere-less fragment. This in turn could result in a different change of the genetic program, that leads to different level of meiotic outcome. We verified that the meiotic phenotypes present in YBT68 were not a result of the 13-base pair deletion in i14 by engineering a strain in which we recreated the wild-type sequence at the i14 locus within the YBT68 strain. This repair did not rescue the brood size or embryonic lethality defects observed in YBT68 (Fig. S2g-h).

Importantly, the MSCI loss we observed in YBT7 was also observed in the YBT68 strain. The relative staining of RNAPII on the HIM-8 track was significantly higher in YBT68 than in wild-type gonads (Fig. 5h-i). These results show that in both of the strains we engineered to have broken X chromosomes, there were defects in MSCI.

### MSCI is aberrant in a strain with reciprocal translocation of chromosomes V and X

We next sought to test whether MSCI is impaired when the X chromosome integrity is compromised using a different technology. Herman et al. previously reported the isolation of SP486, a strain with the mnT10 reciprocal translocation between chromosomes X and V ^75^. Hermaphrodite worms of this strain have a pair of homologous chromosomes with part of chromosome V fused to part of the X chromosome and another pair with the reciprocally fused parts. We stained gonads of SP486 worms with HIM-8 and RNAPII antibodies. Similar to YBT7 and YBT68, in early pachytene nuclei of SP486 the ratio of RNAPII staining between the HIM-8 region and the autosomes was significantly higher than in wild-type gonads (Fig. 6a-b, 130%±20% vs. 58%±3 respectively).

**Figure 6:**
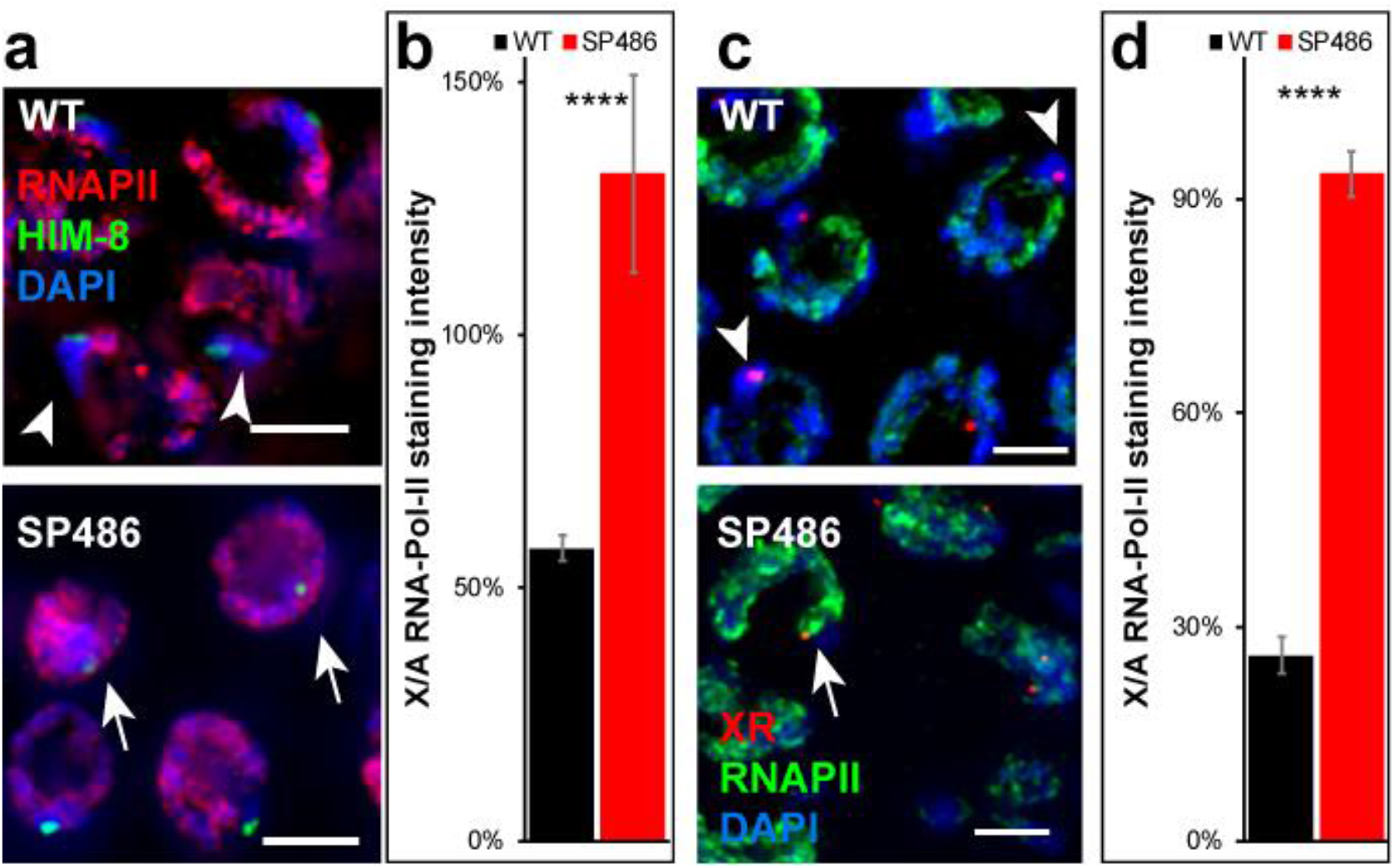
Reciprocal translocation of X and V chromosomes causes defects in MSCI and meiosis. **a,** Early pachytene nuclei stained with DAPI (blue), RNAPII (red), and HIM-8 (green). Scale bar = 3 μM. **c,** Early pachytene nuclei stained with DAPI (blue), RNAPII (green), and FISH probe to the right side of the X (red). Scale bar = 3 μM. **b, d,** Average of the relative RNAPII on the X segment vs an autosome within the same nucleus of **a** and **c** respectively. Arrows: chromosomes with no RNAPII staining. Arrowheads: chromosomes with significant RNAPII staining. n=20. **** *p*<0.0001, Mann-Whitney test.

To the best of our knowledge the break sites that led to the translocation in the SP486 strain have not been fully characterized. It is therefore possible that the region that binds to HIM-8 is only a small part of the X chromosome and most of the X was translocated to a different part of chromosome V and was silenced. To test this possibility, we stained gonads with antibodies against RNAPII and FISH probes directed to the right side of the X chromosome. In wild-type early pachytene nuclei, RNAPII staining was very weak on the right side of the X chromosome compared to that in autosomes (Fig. 6c-d). In SP486 gonads, however, the region stained by the probe directed to the right side of the X chromosome was stained with RNAPII as strongly as were autosomes (Fig. 6c-d, 26%±3% vs. 94%±3% for wild type and SP486, respectively, n=20). These results indicate that MSCI initiation fails when regions of the X chromosome are translocated through natural events. Moreover, these results indicate that MSCI aberrations due to integrity loss are not limited to specific break sites.

### Both synapsed and unsynapsed fragments of the X chromosome are actively transcribed in early meiotic stages

The accepted model for MSCI places the trigger for the inactivation in the unsynapsed region of the sex chromosomes ^17^. In hermaphroditic *C. elegans*, the two X chromosomes synapse, yet undergo MSCI during early oogenesis ^26–29, 54^. MSUC also occurs in hermaphroditic worms ^32^. If MSUC occurs on the X chromosome, we expect that unsynapsed segments of the X will undergo silencing. Alternatively, if unsynapsed segments do not undergo silencing upon sex chromosome integrity loss, the mechanism must differ from MSUC. Due to the nature of pairing in *C. elegans* meiosis, which is required for homolog synapsis, any chromosomal body that harbors a pairing center, pairs and synapses, whereas bodies without paring centers often do not synapse ^52, 76^. The linear fragment of YBT7 contains the pairing center and is therefore expected to pair and synapse, while the circular fragment is expected to stay unsynapsed. If MSCI is a special case of MSUC, then gene expression should be silenced on the unsynapsed fragment. Our data, however, indicate that genes on both the linear and circular fragments of the X chromosome are highly upregulated in YBT7 (Fig. 3). Therefore, either the circular fragment is synapsed, or it escapes MSUC.

To determine the fate of the circular fragment, we stained gonads with antibodies directed against the synaptonemal complex central protein SYP-4, ^77^ and against active RNAPII. In 100% of wild-type early and mid-pachytene nuclei we found a DAPI stained body with a SYP-4 track but without significant RNAPII staining, corresponding to the synapsed and silenced X chromosomes present in wild-type gonads (Fig. 7). In YBT7 gonads, only 5% of mid-early pachytene nuclei had this type of staining combination (*p*<0.00001, Fisher’s exact test, n≥42). In 40% of YBT7 early and mid-pachytene nuclei, all the chromosomes stained with both SYP-4 and RNAPII. These nuclei have either lost the circular fragment, which does not contain a pairing center, or the circular fragment was synapsed (Fig. 7c). In 36% of the YBT7 nuclei, we detected chromosomes stained with RNAPII but not SYP-4 (Fig. 7a-b). These chromosomes were mostly smaller than other chromosomes (Fig. 7b), suggesting that they are fragments of the X chromosome. Since the linear fragment has a pairing center, whereas the circular fragment does not, these unsynapsed active chromosomes are probably the circular fragment. To specifically identity these unsynapsed active chromosomes as the circular fragment, a FISH marker must be added to this staining combination, yet so far we have been unable to make all four stains work simultaneously and reproducibly in the same gonad. Together, these results indicate that in the majority of nuclei from YBT7 gonads, the X fragments were not silenced regardless of whether or not they were synapsed. The presence of the chromosomes that were stained robustly with RNAPII but not SYP-4 show that loss of X chromosome integrity can lead to loss of MSUC. Furthermore, our data suggest that asynapsis of parts of the X do not necessarily lead to silencing. Our findings support the hypothesis that loss of sex chromosome integrity can prevent both meiotic silencing and silencing of unsynapsed chromatin.

**Figure 7:**
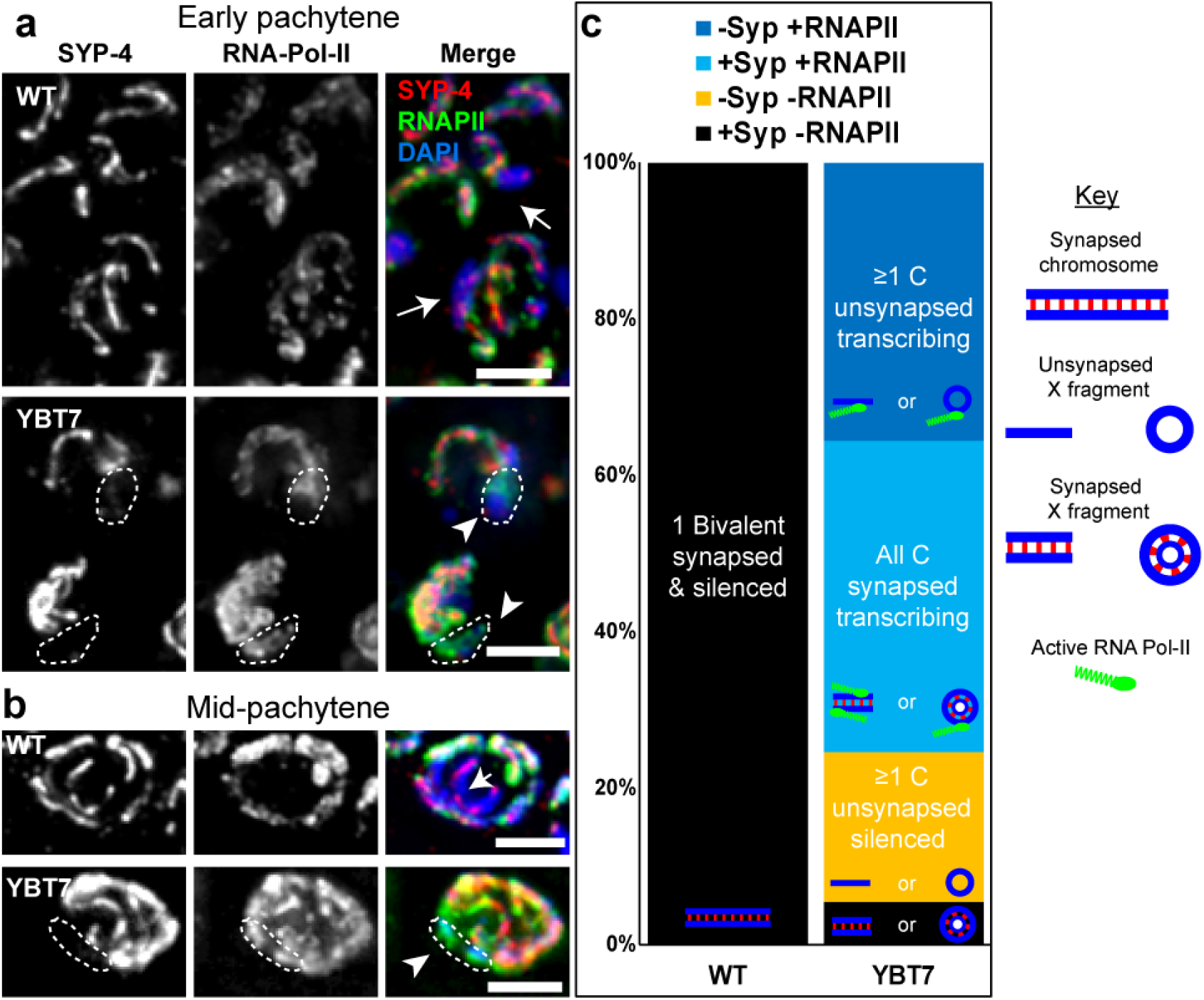
Unsynapsed X segments are transcriptionally active. **a,** Early and **b,** mid pachytene nuclei of wild-type and YBT7 genotypes stained with DAPI (blue), RNAPII (green), SYP-4 (red). Arrows indicate chromosomes with SYP-4 staining without RNAPII staining. Arrowheads indicate chromosomes without SYP-4 staining and with RNAPII staining. Scale bar = 3 μM. **c,** Distribution of the percentages of wild-type and YBT7 early and mid pachytene nuclei with a chromosome stained with SYP-4 and without RNAPII (black), one or more chromosomes stained without SYP-4 and without RNAPII (yellow), all chromosomes stained with SYP-4 and RNAPII (light blue), and one or more chromosome stained without SYP-4 and with RNAPII (blue). Possible interpretations for the different categories are illustrated on the bars: In WT the synapsed chromosome that does not stain with RNAPII is probably chromosome X, and in YBT7 these are the synapsed fragments (the linear, circular, or both). In YBT7, the chromosomes without SYP-4 and RNAPII (yellow in panel c) are fragments of the X which are not synapsed and are silent. Nuclei which only contain chromosomes with SYP-4 and RNAPII (light blue in panel c) have either lost the circular fragment or both fragments are synapsed and active. The chromosomes that stain with RNAPII but not with SYP-4 (blue in panel c) are probably circular fragments that cannot synapse and that escape both MSCI and MSUC.

## Discussion

Sex chromosomes have emerged several times during animal evolution ^78, 79^ as has MSCI ^13–17, 80^. This correlation suggests that evolutionary pressure drives the silencing of sex chromosomes during meiosis. Since MSUC occurs in organisms without sex chromosomes, it is conceivable that when sex chromosomes emerge, they undergo silencing in heterogametic meiocytes simply because they do not synapse (as was suggested previously ^17^). Supporting this model is the finding in various organisms that there is aberrant MSCI and synapsis in sex chromosomes that have undergone translocation with an autosome ^34–37^.

In this work we tested whether superimposed on the synapsis trigger for meiotic silencing, there is another mechanism that depends on sex chromosome integrity. We created strains in which the X chromosome was broken into segments of similar size. Cytological and quantitative transcriptomic evidence supports our conclusion that there are defects in meiotic silencing in these broken segments. Although X-linked genes were highly enriched among the differentially expressed genes in the YBT7 strain, in which the X chromosome is broken into one linear and one circular fragment, a considerable number of autosomal genes were also upregulated. Differential expression of autosomal genes could be due to misregulation of genes on the X chromosome that directly influence autosomal transcription (*e.g.,* transcription factors and chromatin modifiers). Although we found genes that fall into this category (e.g., *lsd-1* and *atf-5*) in the YBT7 strain, we were not able to link these directly to upregulated genes on the autosomes. This suggests that a complex network dysregulated in the mutant results in global alterations in the transcriptome.

We also provide evidence that a reciprocal translocation of the X with an autosome present in SP486 worms leads to defects in MSCI. Kelly et al. reported that in pachytene nuclei of this strain, a region of a chromosome had low levels of H3K4me2 ^26^. We quantified the levels of RNAPII in autosomes and in the two regions of the X chromosome in early pachytene nuclei and found that levels of transcription were significantly increased in the X chromosome regions. This discrepancy could be due to several factors. It is possible that the X-related segments in SP486 gonads are not enriched with histone modifications corelated with transcription, even though transcription is occurring. Alternatively, since Kelly et al. did not use cytological markers to identify chromosomes, it is possible that the fragments they observed were not part of the X, but rather an unsynapsed part of chromosome V. We noticed high variability of the RNAPII staining levels on the X segments in these nuclei, and only reached our conclusion following careful quantification. This type of variability in MSCI markers of translocated X segments is not limited to *C. elegans*. For example, Turner et al. used γH2AX as a marker for MSCI initiation in mouse testes with a reciprocal translocation of chromosomes 16 and X and found that of 72 pachytene spermatocyte nuclei with synapsed X^16^ scored, in 38 nuclei the X part was stained with γH2AX, and in 34 it was not ^37^. Thus, in the system studied by Turner et al. the X silencing occurs in about 50% of synapsed X chromosome regions. Similarly, Mary et al. reported that in a boar with translocation of chromosomes 13 and Y, about 50% of the Y and X chromosomes showed no γH2AX signal in pachytene spermatocytes nuclei ^38^. Similar levels were reported by Barasc et al. in a boar with translocation of chromosomes 1 and Y ^35^. These reports indicate that in both worms and mammals, reciprocal translocation of a sex chromosome to an autosome incompletely perturbs MSCI. The variability in the silencing observed in translocations involving sex chromosomes and autosomes could also arise due to dynamics of epigenetic modifications.

Our finding that in some cases the unsynapsed segments of the X chromosomes were not silenced was surprising given that the accepted model views the meiotic silencing of sex chromosomes as a special case of the silencing of unsynapsed chromatin. This uncoupling of synapsis and silencing of chromatin derived from sex chromosomes may be due to an epigenetic mechanism. Compared to YBT7, the percentages of nuclei with unsynapsed and actively transcribed X chromosome fragments were much lower in the other strains that lost the X integrity (data not shown), and we presume that in these strains the original “identity” of the chromatin was not faithfully maintained. Another possibility is that all the fragments were synapsed. It will be important to determine whether this uncoupling of synapsis and expression is unique to *C. elegans* hermaphrodites in which the X chromosomes do synapse yet still undergo silencing. Several lines of evidence suggest that this feature is evolutionarily conserved. First, silencing of unsynapsed chromatin has been observed in *C. elegans* ^32^, so the basic mechanism of MSUC exists. Second, disruption of sex chromosome integrity in mammals leads to silencing of synapsed parts of autosomes and sex chromosomes in pachytene ^34-36^. Third, in early pachytene nuclei of XO mice, the unsynapsed X chromosome is marked by γH2AX yet is transcribed ^30^. Taken together these reports imply that sex chromosomes undergo silencing in heterogametic wild-type meiocytes not solely due to their unsynapsed state.

Considering these previous reports and the findings we report here, we propose the following model: Under normal conditions MSCI is activated on complete sex chromosomes in heterogametic meiocytes, as well as in early stages of hermaphrodite worms. Lack of synapsis in autosomes leads to silencing, whereas aberrant synapsis of sex chromosomes cancels their silencing. When sex chromosomes break, at least in *C. elegans* hermaphrodites, another mechanism is activated, and the silencing of sex chromosomes is perturbed. This mechanism can in some meiocytes override MSUC, and the unsynapsed fragments are transcribed.

What evolutionary drive links silencing of sex chromosomes to their integrity? One possible answer comes from inherent problems with DNA repair of heterogametic chromosomes during meiosis. As interhomolog recombination is the preferred repair pathway of meiotic breaks, heterogametic chromosomes are at risk of aberrant repair and breakage. Therefore, loss of MSCI when sex chromosome integrity is compromised may simply be a safety mechanism that eliminates meiocytes in which the sex chromosomes are fragmented. The integrity loss will in turn disrupt MSCI, which will lead to gametogenesis failure and apoptosis. Alternatively, flexibility in the dichotomic chromosome state between silent sex chromosomes and active autosomal chromosomes may allow formation and disappearance of sex chromosomes with evolutionary progression.

Like initiation of silencing, other meiotic processes are also executed differently on the X chromosome than on autosomes in hermaphroditic worms ^81–86^. These differences suggest that the X chromosome is marked differently than the autosomes, and, indeed, previous reports indicate that the X chromosomes are enriched with different histone modifications than are autosomes ^26, 32, 81, 82, 87^. Thus, an epigenetic mechanism may regulate silencing and its dependence on integrity. Our results suggest the possibility that the silencing initiation depends on an output of sex chromosome that assess integrity. One potential regulator is the synaptonemal complex. Axis length in *C. elegans* appears to regulate DNA double-strand break formation and crossover interference ^88, 89^, and axis proteins are in close contact with chromatin. Additional studies should test these possibilities to determine if these regulators connect X chromosome integrity to meiotic silencing.

## Methods

### Strains and alleles

All strains were cultured under standard conditions at 20 °C unless specified otherwise ^90^. The N2 Bristol strain was utilized as the wild-type background. Worms were grown on NGM plates with *Escherichia coli* OP50 ^90^. All experiments were conducted using adult hermaphrodites 20-24 h after the L4 stage. The following mutations and chromosome rearrangements were used: SP486: mnT10 (V;X) ^75^. Strains engineered in this work (see below): YBT7: *deg-1(huj32) linc-20(huj2) hujCf1*, YBT68: *deg-1(huj33) linc-20(huj29) hujCf2,* YBT54: *linc-20(huj21),* YBT67: *deg-1(huj28),* YBT75: *linc-20(huj29)* and YBT72: *deg-1(huj32) linc-20(huj2).*

### Generation of strains by CRISPR-Cas9 genome engineering

To generate the YBT7 strain, we used the procedure described in ^91^ with the modifications detailed previously ^50^. The gRNA sequences are given in Table S3. Worms were isolated based on PCR analysis of targeted loci, and broken chromosomes were identified via Nanopore sequencing as described below. The YBT7 strain carries the deletions *deg-1(huj32)* X:7769748-7772344 and *linc-20(huj2)* X:16508000-16509415 and X:16511093-16513798. A fusion of the region X:~7772k to X:~16511k and fusion of the segment left of X:7774k to segment right of 16507k were confirmed by Sanger sequencing. Illumina DNA sequencing indicated that there are other structural aberrations and mutations, but these were not confirmed by Sanger sequencing.

All other strains were generated using the protocol described previously ^73^ with the modifications detailed previously ^50^. YBT68: *deg-1(huj33) linc-20(huj29) hujCf2* was generated using crispr RNAs (crRNAs) listed in Table S3, isolated in a similar strategy to YBT7, and outcrossed five times. This strain carries the deletions *deg-1(huj33)* X:7769768-X:7769760, *linc-20(huj29)* X:16507326-16508724 and X:16509401-16510374. Nanopore sequencing suggested that the right and the left halves of the X are not connected, and cytological observation supported this hypothesis. Illumina DNA sequencing suggested the presence of other structural aberrations and mutations, but these were not confirmed by Sanger sequencing.

YBT54: *linc-2(huj21)* was generated using crRNAs listed in Table S3 and was outcrossed five times. This strain carries the deletion *linc-20(huj21)* X:16507934-16509755. YBT67: *deg-1(huj28)* was engineered using crRNAs listed in Table S3. It has a 4-base out-of-frame deletion at position 297 of the first exon. YBT75: *linc-20(huj29)* was engineered using appropriate crRNAs and single-stranded oligodeoxynucleotides (ssODNs) (Table S3). These were injected into YBT68 worms, and repair of the *deg-1* genotype was verified by sequencing.

YBT72 was engineered through three CRISPR engineering steps as follows: Wild-type worms were engineered using crRNAs “homologous lincs 5’ crRNA” and “homologous lincs 3’ crRNAs” together with the ssODN “Linc-20 del1-ssODN” and “Linc-20 del2-ssODN”. The strain with *linc-20* (*huj21)* was identified by PCR and verified by sequencing. After one outcross with the wild-type strain, worms were further engineered with “Linc-20 YBT7 del-2 5’ crRNA” and “Linc-20 YBT7 del-2 3’ crRNA” together with “Linc-20 del2-ssODN”, and worms with *linc-20(huj2)* were identified by PCR and verified by sequencing and outcrossed once with the wild-type strain. A strain with *deg-1(huj32)* was engineered by injecting wild-type worms with crRNAs “Deg-1 YBT7 del-5’ crRNA” and “Deg-1 YBT7 del-3’ crRNA” and ssODN “Deg-1 del-ssODN” (Table S3) and outcrossed once. Worms with *deg-1(huj32)* were crossed with worms with *huj(2)* to establish YBT72. All the engineered mutations were verified by Sanger sequencing.

### Cytological analysis and immunostaining

DAPI and immunostaining of dissected gonads were carried out as described ^61, 92^. Worms were permeabilized on Superfrost+ slides for 2 min with methanol at −20 °C and fixed for 30 min in 4% paraformaldehyde in phosphate-buffered saline (PBS). Staining with 500 ng/ml DAPI was carried out for 10 min, followed by destaining in PBS containing 0.1% Tween 20 (PBST). Slides were mounted with Vectashield anti-fading medium (Vector Laboratories). Primary antibodies were used at the following dilutions: rabbit anti-SYP-4 (1:200, a kind gift from S. Smolikove, The University of Iowa), rabbit anti-HIM-8 (Novus Biological, 1:2000), rat anti-HIM-8 (1:100, a kind gift from A. Dernburg, University of California, Berkeley), rabbit anti-H3K27me3 (Millipore, 1:1000), rabbit anti-H3K4me2 (Millipore, 1:1000), and mouse anti-pSer2 RNAPII (Diagenode, 1:1000). The secondary antibodies used were Cy2-donkey anti-rabbit, Cy3-donkey anti-goat, Cy2-goat anti-rat, Cy2-goat anti-mouse, Cy3-goat anti-mouse (Jackson ImmunoResearch Laboratories).

### DNA FISH

Probes were made from cosmids provided by the *C. elegans* sequencing consortium at the Sanger Centre. Cosmid DNAs that harbor 30-40 kb of sequence around the chosen genomic target were labeled after linearization by nick translation using cy3dUTP (GE Healthcare) as described ^93^. For the region left of *linc-20* we used C09G1, and for the right side of the X chromosome we used T27B1.

Worms were transferred to a 15-μL drop of egg buffer (118 mM NaCl, 48 mM KCl, 2 mM CaCl_2_, 2 mM MgCl_2_, 5 mM HEPES [pH 7.4]) ^94^, containing 15 mM NaN_3_ and 0.1% Tween-20 on a 22×22 mm coverslip. Gonads were dissected and fixed in 3.7% formaldehyde in PBST. A SuperFrost Plus slide (ThermoFisher Scientific) was placed on the coverslip, then frozen on an aluminum block immersed in liquid nitrogen. The coverslip was cracked off and slides were transferred to methanol at −20 °C for 30 min. Slides were washed in 2X SSCT (3 M NaCl, 0.3 M sodium citrate, pH 7, 0.1% Tween20), 25% formamide/2X SSCT and incubated for 4 h at 37 °C in 50% formamide/2X SSCT in a humid chamber. Slides were prehybridized on a heat block at 93 °C for 90 s in hybridization solution (50% formamide, 3× SSC, 10% dextran sulfate) containing 1 μl of the labeled probe. Slides were hybridized overnight in a humid chamber at 37 °C. After washing with 2X SSCT, the slides were either directly labeled with DAPI and mounted in Vectashield solution for visualization or were blocked 30 min at room temperature in 1% BSA before antibody labeling.

### Imaging and microscopy

Z-stack 3D images shown in Figure 2 were acquired at 0.3 μm increments using an Olympus FV1000 Inverted Confocal IX81 Microscope and FV10-ASW 3.1 Software (Olympus). All other images were acquired using the Olympus IX83 fluorescence microscope system. Optical Z-sections were collected at 0.30- or 0.60-μm increments with the Hamamatsu Orca Flash 4.0 v3 and CellSens Dimension imaging software (Olympus). Pictures were deconvolved using AutoQuant X3 (Media Cybernetics).

### Progeny and embryonic lethality quantification

Brood sizes and embryonic lethality were determined by placing individual L4 worms on seeded NGM plates, transferring each worm to a new plate every 24 h, and counting embryos and hatched progeny during a 3-day period.

### Analysis of synapsis and expression interactions

Early and mid-pachytene nuclei stained with DAPI, anti-SYP-4, and anti pSer2 RNAPII were captured at 0.3 μM optic Z intervals. Nuclei were binned into one of four categories: 1) at least one chromosome positive for RNAPII and negative for SYP-4, 2) all chromosomes positive for both markers, 3) all SYP-4-negative chromosomes are also negative for RNAPII, and 4) all chromosomes are SYP-4 positive and at least one is RNAPII negative.

### RNA-seq

Gonads were manually dissected from worms at 24 h post L4 and immediately placed in Eppendorf tubes with Trizol reagent. After several freeze-crack cycles in liquid nitrogen, total RNA was extracted using Zymo Research Direct-zol RNA miniprep plus kit. Synthesis of first strand was done on 10 μg of total RNA using ThermoFisher SuperScript III Reverse Transcriptase with the following primer that includes the T7 promotor, a unique molecular identifier, UMI and polyT: 5’-CGATGACGTAATACGACTCACTATAGGGATACCACCATGGCTCTTTC CCTACACGACGCTCTTCCGATCTNNNNNNNNNNTTTTTTTTTTTTTTTTTTTVN-3’. Removal of excess primers was done using New England Biolabs Exonuclease I and ThermoFisher FastDigest HinfI in provided buffers; samples were incubated 45 min at 37 °C and then 10 min at 80 °C. The product was purified using Beckman AMPure XP magnetic beads, eluted in 14.5 μL of 10 mM Tris, followed by second-strand cDNA synthesis using New England Biolabs NEBNext Ultra II Non-Directional RNA Second Strand Synthesis Module. Samples were concentrated to 8 μL, and then the product was transcribed with the New England Biolabs HiScribe T7 High Yield RNA Synthesis Kit. RNA was purified using AMPure XP beads and eluted in 20 μL of 10 mM Tris. A 9-μL aliquot of RNA was fragmented using Invitrogen RNA Fragmentation Reagents kit for 3 min. Fragments were purified using AMPure XP beads and eluted in 11 μL Tris. Synthesis of first-strand cDNA and was performed using ThermoFisher SuperScript III Reverse Transcriptase using PvG748 primer 5’-AGACGTGTGCTCTTCCGATCTNNNNNN-3’. After purification using AMPure XP beads and elution with 12.5 μL 10 mM Tris, libraries were amplified using Kapa Biosystems HiFi HotStart ReadyMix, with 2p fixed primers (2p Fixed, 5’-AATGATACGGCGACCACCGAGATCTACACTCTTTCCCTACACGACGCTCTTCCGATCT-3’ and 2p Fixed +barcode, 5’-CAAGCAGAAGACGGCATACGAGATNNNNNNNNGTGACTGGAGTTCAGACGTGTGCTCTTCCGATCT-3’). The product was purified using AMPure XP beads and eluted in 32 μL of doubly distilled water. Deep sequencing was carried out on an Illumina NextSeq following the manufacturer’s protocols; >38 million reads were generated for each sample.

### Differential expression analysis

Raw reads were trimmed off low quality and technical bases. Cutadapt, version 1.12, with parameters -O 1, -m 15 and --use-reads-wildcards. Reads with overall low quality were removed using fastq_quality_filter, FASTX version 0.0.14, with parameters -q 20 and -p 90. Processed reads were aligned to the *C. elegans* genome version WBcel235 using TopHat2, version 2.1.1. Alignment allowed 2 mismatches and 5-base gaps, and used gene annotations from Ensembl release 36. Raw counts per gene were calculated with htseq-count, version 0.6.0, using default parameters.

Normalization and differential expression were calculated with the R package DESeq2, version 1.12.4. Calculations were done for genes with at least 3 raw counts using default parameters. Genes were taken as differentially expressed if their baseMean was above 5 and if the absolute maximum likelihood estimate of the fold change (without shrinkage, lfcMLE) was greater than 5/baseMean^0.5 + 1. This baseMean-dependent threshold for the change in expression required at least 2-fold change in expression for highly expressed genes, and the requirement becomes stricter as the level expression becomes lower. The MA plot is illustrated in Fig. S1.

### Nanopore DNA sequencing

Worms were washed from NGM plates with M9 buffer, and young adult worms were isolated on a 60% sucrose bed. Worms were then washed in M9 buffer and frozen in liquid nitrogen. DNA was isolated using Zymo Research Quick-DNA Miniprep kit.

Genomic DNA was barcoded without fragmentation using Oxford Nanopore Technologies EXP-NBD103, SQK-LSK108/9 according to the vendor’s instructions. Approximately 260 ng DNA of each strain were loaded two in a cell couples on one MinION flowcell (Oxford Nanopore Technologies), and sequencing was performed using GridION device and MinKnow software for 48 h.

using command line Guppy (version 3.4.4), and reads were quality filtered using NanoFilt (version 2.6.0, parameters ‘-q 5 -l 100 --headcrop 40’). Filtered reads were aligned to the *C. elegans* genome (WBcel235) using minimap2 (version 2.17^95^). A combination of three tools were used for identification of structural variations: sniffles (version 1.0.11, ^96^), NanoSV (version 1.2.3, ^97^), and SVIM (version 1.2.0, ^98^). Copy number variations were identified using the R package QDNAseq ^99^.

### Illumina DNA sequencing

DNA was extracted from 25 μL of packed young adult worms using Gentra Puregene Tissue Kit (Qiagen) according to vendor protocol for *C. elegans*. For each sample, 1000 ng of DNA was sheared using the Covaris E220X sonicator. End repair was performed in an 80-μL reaction at 20 °C for 30 min. After purification using AMPURE XP beads in a ratio of 0.75X beads to DNA volume, A bases were added to both 3’ ends followed by adapter ligation in a final concentration of 0.125 μM. A solid-phase reversible immobilization (SPRI) bead cleanup in a ratio of 0.75x beads to DNA volume was performed, followed by eight PCR cycles using 2X KAPA HiFi ready mix in a total volume of 25 μL with the following program: 2 min at 98 °C, 8 cycles of 20 s at 98 °C, 30 s at 55 °C, 60 s at 72 °C, and 10 min at 72 °C.

Libraries were evaluated by Qubit and TapeStation. Sequencing libraries were constructed with barcodes to allow multiplexing of four samples on one lane. Between 38-45 million paired-end 150-bp reads were sequenced on Illumina Nextseq 500 instrument Mid output 300 cycles kit.

Reads were mapped to the *C. elegans* genome (Ensembl’s WBcel235) using bwa-0.7.5a ^100^ mem algorithm and then deduplicated using Picard tools v.2.8.1. Variant calling was done with GATK’s Haplotype caller v3.7 ^101^. Variants were filtered with the following values for single-nucleotide polymorphisms and indels, respectively: QD<2.0, FS>60.0, MQ<40.0, HaplotypeScore>13.0, MQRankSum<-12.5, ReadPosRankSum<-8.0 and QD<2.0, FS>200.0 and ReadPosRankSum<-20.0. Variants were then annotated with Ensembl’s Variant Effect Predictor v.83 ^102^.

### Measuring distances between chromosome markers

To measure the spatial distance between chromosomal makers, mid-pachytene nuclei positively stained for DAPI and HIM-8 and with the FISH probe were completely captured at 0.3 μm Z increments. The distance between the HIM-8 foci and the FISH probe was measured using ImageJ. Significance was estimated via the Mann-Whitney test.

### Relative staining intensity of expression markers of X chromosome vs. autosomes

To measure the level of expression marker stainin<<g, early pachytene nuclei positively stained for DAPI and H3K27me3, H3K4me2, or pSer2 RNAPII were captured. The staining level on the X chromosome (marked by either HIM-8 or the FISH probe directed to the right side of the X) was measured in ImageJ, as well on another chromosome within the same nucleus. The ratio for each nucleus was calculated and averaged across all nuclei.

## Availability of data and materials

Strains and plasmids are available upon request. Table S3 contains detailed descriptions of all primers used for genome engineering and genotyping.

## Acknowledgments

We thank the Caenorhabditis Genetics Center for kindly providing strains. We thank Abby Dernburg for the HIM-8 antibody and Sarit Smolikove for the SYP-4 antibody. We thank Yuval Nevo and the team at the Info-CORE, Bioinformatics Unit of the I-CORE Computation Center, The Hebrew University and Hadassah Medical Center, Jerusalem, Israel, for the differential expression analysis. We thank Michal Bronstein and the team at the Center for Genomic Technologies of the Alexander Silberman Life Sciences Institute for their help with Nanopore sequencing. We thank Michael Gershovis and the team of Israel National Center for Personalized Medicine for their help in Illumina DNA genomic sequencing. This work was supported by the European Research Council (ERC, #715260 SC-EpiCode), the Israeli Center of Research Excellence (I-CORE) program, the Israel Science Foundation (ISF, #1618/16), and Azriely Foundation Scholar Program for Distinguished Junior Faculty to O.R. and by the Israel Science Foundation (grants #1283/15 and #2090/15) to Y.B.T.

## Supplementary

**File S1: Nanopore data demonstrating fragmentation of the X chromosomes in YBT7 and YBT68 strains.**

**Figure S1: Differential gene expression in YBT7 and wild-type gonads.** Expression differences in YBT7 vs. wild-type gonads for each gene plotted against the average expression. The X axis is the DESeq2-calculated baseMean, and the Y axis is the DESeq2-calculated lfcMLE (log2 of the fold-change maximum likelihood estimate). Orange dots indicate differentially expressed genes, other genes are indicated as black dots. The Y axis was limited to a range between −6 and 6; triangles represent genes with lfcMLE value beyond this range.

**Figure S2: Meiotic defects in YBT7 are the result of the chromosome breaks and not local changes**. **a,** Average progeny brood size per worm and, **b,** embryonic lethality of wild type vs *deg-1(huj28*) (disruption within *deg-1,* n≥8). **c,** Average progeny brood size per worm and, **d,** embryonic lethality of wild type vs *linc-20(huj21*) (full deletion of *linc-20,* n=7). **e,** Average progeny brood size per worm and, **f,** embryonic lethality of wild type vs YBT72 (deletions within *deg-1* and *linc-20,* identical to YBT7, n=10). **g,** Average progeny brood size per worm and, **h,**embryonic lethality of wild type vs YBT75 (YBT68 with reconstructed WT *deg-1*, n≥6). * *p*<0.05, ** *p*<0.01, N.S. indicates not significant, Mann-Whitney test.

**Table S1: Differential expression in YBT7 versus wild-type gonads.** Columns include general gene details (A-F), raw counts for each strain (G and H), normalized counts (I and J), the DESeq2 calculated baseMean (K), the DESeq2 calculated lfcMLE (L), and a column for the final decision of differential expression (M). Genes with less than 3 counts are indicated with question marks in columns I to M. There is a question mark in the general gene details when information was missing from the database. The final decision column (M) has the value ‘THR’ for differentially expressed genes (i.e., those with a baseMean greater than 5 and a lfcMLE greater than 5/baseMean^0.5 + 1) and a value ‘FALSE’ for all other genes.

**Table S2: Point mutations in YBT7 and YBT68.** Homozygous mutations detected by Illumina sequencing in YBT7 and YBT68 but not in the parental wild-type strain.

**Table S3: List of gRNAs, crRNAs, and ssODNs used for genome engineering.**

## Notes

### Competing Interest Statement

The authors have declared no competing interest.

